# DELPHI: accurate deep ensemble model for protein interaction sites prediction

**DOI:** 10.1101/2020.01.31.929570

**Authors:** Yiwei Li, Lucian Ilie

## Abstract

**Motivation:** Proteins usually perform their functions by interacting with other proteins, which is why accurately predicting protein-protein interaction (PPI) binding sites is a fundamental problem. Experimental methods are slow and expensive. Therefore, great efforts are being made towards increasing the performance of computational methods.

**Results:** We propose DELPHI (DEep Learning Prediction of Highly probable protein Interaction sites), a new sequence-based deep learning suite for PPI binding sites prediction. DELPHI has an ensemble structure with data augmentation and it employs novel features in addition to existing ones. We comprehensively compare DELPHI to nine state-of-the-art programs on five datasets and show that it is more accurate.

**Availability:** The trained model, source code for training, predicting, and data processing are freely available at https://github.com/lucian-ilie/DELPHI. All datasets used in this study can be downloaded at http://www.csd.uwo.ca/~ilie/DELPHI/.

## Introduction

Protein-protein interactions (PPI) play a key role in many cellular processes such as signal transduction, transport and metabolism [53]. Proteins interact by forming chemical bonds with other proteins. The bonding amino acid residues are protein-protein interaction binding sites. Detecting PPI binding sites helps understand cell regulatory mechanisms, locating drug target, predicting protein functions [8]. Databases like PDB [7] store protein binding sites information deriving from the 3D structure of each protein, but the available protein structures are limited. Experimental methods such as two-hybrid assay and affinity systems are usually time and labour intensive [39]. Computational methods are needed to bridge the gap, and many have been developed [?, 3,4,9–11,15,18,20,22,23,25,28–31,33,34,40,44,45,48,49,51,52,54]. Out of the above mentioned twenty six computational methods, all but one are machine learning based. Computational methods can be classified into three categories, sequenced based, structure based, and combined. Among them, sequence-based approaches are usually faster and cheaper. They are also more universal because comparing to sequence information, structure information is still limited.

Machine learning methods use feature groups to represent each protein sequence. Widely used features such as position-specific scoring matrix (PSSM), evolutionary conservation (ECO), putative relative solvent accessibility (RSA) have been assessed in [55]. High-scoring segment pair (HSP) has been used in previous methods for PPI prediction [27]. One-hot vectors [51,52] and amino acid embedding [5,6,19] have also been empirically explored to represent protein sequences.

The learning structure is crucial to PPI binding sites classification problems. Previously explored architectures include random forest [?,45], SVM [45], logistic regression [54], Bayes classifier [33], artificial neural networks [40]. Recently, convolutional neural network (CNN) [51] and recurrent neural network (RNN) [52] have also been applied to solve this problem.

We introduce a new sequence-based PPI binding sites prediction method, DELPHI (DEep Learning Prediction of Highly probable protein Interaction sites), that combines a CNN and a RNN structure. It uses twelve feature groups to represent protein sequences including three novel features, high-scoring segment pair (HSP), position information, and a reduced 3-mer amino acid embedding (ProtVec1D).

We have comprehensively compared DELPHI with nine state-of-the-art programs on five datasets. DELPHI provides the best predictions in all metrics. The contributions of the DELPHI study are as follows. First, a novel fine tuned ensemble model combing CNN and RNN is constructed in which three features are used the first time in this problem. Second, a many-to-one structure, that not only serves as a data augmentation technique but also improves the prediction performance, is applied. Third, a data processing and feature construction suite, including new features, is provided, aiming to alleviating the difficulty of tedious feature computation by the users.

## Materials and Methods

### Datasets

#### Training and validation datasets

A large, high quality dataset was provided in [55]. In this dataset, Uniprot sequences are annotated with Protein, DNA, RNA, and small ligands binding information at the residue level. We further processed this dataset as follows. First, we kept only the sequences with protein-protein binding information to focus on protein-protein binding. Then we removed any sequences from training dataset sharing more than 25% similarities, as measured by PSI-CD-hit [17,26], with any sequences in testing datasets. It is well acknowledged that similar sequences between training and testing datasets negatively affect the generalization of the evaluated performance of a machine learning model. Also, proteins with higher levels of similarity can be accurately predicted by the alignment-based methods [53]. The similarity threshold is picked differently by different programs ranging from 25% to 50%. We picked the strictest value of 25% to match to one of the closest competing programs, SCRIBER [54], for a fair comparison. We used PSI-CD-HIT because it is fast, accurate and well maintained in the CD-HIT suite. Also, it is able to cluster sequences with similarity at low as 25%, whereas CD-HIT works only down to 40%. Finally, we ran PSI-CD-hit again on the rest of the training protein sequences so no sequences shared more than 25% similarities among training data. This ensures the training data is as diverse as possible. A dataset of 9,982 protein sequences was constructed. From it, we randomly pick eight ninth (8,872) as the training dataset and one ninth (1,110) as the validation dataset.

#### Testing datasets

Five datasets are used in the comparative assessment. Four of them are publicly available dataset from previous studies: Dset_186 [33], Dset_72 [21], Dset_164 [13], and Dset_448 [54]. The first three have been widely used and explored by previous studies while Dset_448 is more recent. The raw data of Dset_448 was from the BioLip database [50] where binding sites are defined if the distance between an atom of a residues and an atom of a given protein partner j0.5 Å plus the sum of the Van der Waals radii of the two atoms. The raw data was further processed by the authors of [54], removing protein fragments, mapping BioLip sequences to UniProt sequences, clustering so that no similarities above 25% is shared within the dataset. The number of sequences, binding, non-binding, and the ratio of binding residues are shown in Table 1. The training dataset of DLPred has some overlaps with Dset_448, therefore we removed 93 proteins in Dset_448 that share more than 40% similarities with DLPred training data and formed a reduced testing dataset of 355 proteins (Dset_355). According to the PSI-CD-HIT results, the training datasets of DLPred and SCRIBER still contain some proteins that share more than 25% similarity to Dset_186, Dset_72, and Dset_164. However, we kept these testing datasets untouched because otherwise we would have to remove too many proteins from them. The training datasets of the competing programs can not be changed because the models are pre-trained. Note that all testing data share less than 25% similarity to the training dataset of DELPHI.

**Table 1.** The datasets used for training, validation, and the comparative testing. The first column are the dataset names. The second column contains the number of proteins in each dataset. The third, fourth, and fifth columns represent the total number of residue, the number of binding, and the number of non-binding residues in each dataset. The last column represents the percentage of the binding residues out of total.

| Dataset | Proteins | Residues |  |  | % binding<br>out of total |
| --- | --- | --- | --- | --- | --- |
|  |  | total | binding | non-binding |  |
| Dset_448 | 448 | 116,500 | 15,810 | 100,690 | 13.57% |
| Dset_355 | 355 | 95,940 | 11,467 | 84,473 | 11.95% |
| Dset_186 | 186 | 36,219 | 5,517 | 30,702 | 15.23% |
| Dset_72 | 72 | 18,140 | 1,923 | 16,217 | 10.60% |
| Dset_164 | 164 | 33,681 | 6,096 | 27,585 | 18.10% |
| Training + validation | 9,982 | 4,254,198 | 427,687 | 3,826,511 | 10.05% |

### Input features

DELPHI uses 12 features groups, shown in Table 2, including high-scoring segment pair (HSP), a variation of 3-mer amino acid embedding (ProtVec1D), position information, position-specific scoring matrix (PSSM), evolutionary conservation (ECO), putative relative solvent accessibility (RSA), relative amino acid propensity (RAA), putative protein-binding disorder, hydropathy index, physicochemical characteristics, physical properties, and PKx. Each input is represented by a 39 dimensional feature vector profile. To the best of our knowledge, this study is the first time that HSP and ProtVec1D are being used in binding sites classification problems. The computation of each of these two new features is described next. High-scoring segment pair (HSP): An HSP is a pair of similar sub-sequences between two proteins. The similarities between two sub-sequence of the same length are measured by scoring matrices such as PAM and BLOSUM. SPRINT [27] is used for computing all HSPs as it detects similarities fast and accurately among all proteins in training and testing. After obtaining the HSPs, the score for the *i*th residue, *P*[*i*], of a testing protein *P*, denoted HSP_score_(*P*[*i*]), is calculated as follows. Assume we have an HSP, (*u,v*), between *P* and a training protein *Q* such that *u* covers the residue *P* [*i*], that is, position *i* in *P* is within the range covered by *u*. Let *j* be the position in *Q* that corresponds to *i*, that is, the distance in *P* from the beginning of *u* to *i* is the same as the distance in *Q* from the beginning of *v* to *j*. If *Q*[*j*] is a known interacting residue, then we add the PAM120 score between *P*[*i*] and *Q*[*j*] to the HSP score of *P*[*i*]:

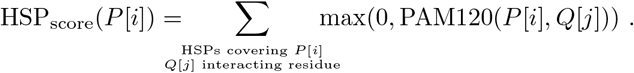

**Table 2.** Feature groups used by DELPHI. The first column indicates the name of each feature. The second column describes the program used to obtain the feature. “Load” means the value for a specific amino acid is fixed, and it is loaded in DELPHI program. “Compute” means DELPHI further computes the feature. The last column shows the dimension of each feature group.

| Feature | Program | Dimension |
| --- | --- | --- |
| High-scoring segment pair (HSP) | SPRINT and compute | 1 |
| 3-mer amino acid embedding (ProtVec1D) | Load and compute | 1 |
| Position information | Compute | 1 |
| Position-specific scoring matrix (PSSM) | Psi-Blast | 20 |
| Evolutionary conservation (ECO) | Hhblits | 1 |
| Putative relative solvent accessibility (RSA) | ASAquick | 1 |
| Relative amino acid propensity (RAA) | Load | 1 |
| Putative protein-binding disorder | ANCHOR | 1 |
| Hydropathy index | Load | 1 |
| Physicochemical characteristics | Load | 3 |
| Physical properties | Load | 7 |
| PKx | Load | 1 |

The 3-mer amino acid embedding (ProtVec1D): We developed this feature based on ProtVec [6]. ProtVec uses word2vec [32] to construct a one hundred dimensional embedding for each amino acid 3-mer. It is shown in [6] that ProtVec can be applied to problems such as protein family classification, protein visualization, structure prediction, disordered protein identification, and protein-protein interaction prediction. Since using the ProtVec embedding in our program slows down significantly the deep learning model, especially during training, we replaced the one hundred dimensional vector by one dimensional value, which is the sum of the one hundred components; we call this ProtVec1D. According to our tests, ProtVec1D achieves, in connection with the other feaures, the same prediction performance as ProtVec.

Position information: In natural language processing tasks, position information is shown useful. The popular network Bert [?] utilizes this information to guide its translation process. It is also shown by DeepPPISP [51] that the global information of a protein helps the prediction of interfaces. Inspired by the two networks, we use the position information of each amino acid as an input feature hoping that it provides certain global information of a protein. The position of an amino acid in a protein is in the range of 1 to the length of the protein. Then the position is divided by protein’s length so that the value is between 0 to 1.

Position-specific scoring matrix (PSSM): PSSM matrices are widely used in protein interaction related problems. They contain the evolutionary conservation of each amino acid position by aligning an input sequence with protein databases. The PSSM matrices are computed using PSI-Blast [2] with the expectation value (E-value) set to 0.001 and the number of iterations set to 3. PSI-Blast performs multiple alignment on each input sequence against the non-redundant database.

Evolutionary conservation (ECO): ECO also contains evolutionary conservation, but in a more compact way. To compute the ECO score, the faster multiple alignment tool HHBlits [38] is run against the non-redundant Uniprot20 database with default parameters. The one dimensional conservation value is computed using the formula described in [55].

Putative relative solvent accessibility (RSA): The solvent accessibility is predicted using ASAquick [16]. The values are obtained in the from rasaq.pred file in each output directory.

Relative amino acid propensity (RAA): The AA propensity for binding is quantified as relative difference in abundance of a given amino acid type between binding residues and the corresponding non-binding residues located on the protein surface. The RAA for each amino acid type is computed in [55] by using the program of [43].

Putative protein-binding disorder: The putative protein-binding disorder is computed using the ANCHOR program [14].

Hydropathy index: Hydrophobicity scales is experimentally determined transfer free energies for each amino acid. It contains energetics information of protein-bilayer interactions [47]. The values are computed in [24].

Physicochemical characteristics: For each protein, this includes three features: the number of atoms, electrostatic charges and potential hydrogen bonds for each amino acid. They are taken from [52].

Physical properties: We use a 7-dimensional property of each amino acid type. They are a steric parameter (graph-shape index), polarizability, volume (normalized van der Waals volume), hydrophobicity, isoelectric point, helix probability and sheet probability. The pre-computed values are taken from [52].

PKx: This is the negative of the logarithm of the dissociation constant for any other group in the molecule. The values for each amino acid type is taken from [52].

After computing all the feature vectors, the values in in each row vector are normalized to a number between 0 to 1 using formula (1) where *v* is the original feature value, and max and min are the biggest and smallest value observed in the training dataset, resp. This is to ensure each feature group are of the same numeric scale and help the model converge better:

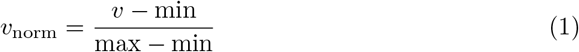

## 0.1 Model architecture

DELPHI has an architecture that is inspired by ensemble learning. The intuition of the design is that different components of the model capture different information, and another deep neural network is trained to only select the most useful ones. As shown in Fig. 1, the model consists of three parts, a convolutional neural network (CNN) component, an recurrent neural network (RNN) component, and an ensemble component. The core layers of the CNN and RNN components are convolution and bidirectional gated recurrent units (GRU) layers. The ensemble model decodes the output of the first two components.

**Figure 1.**
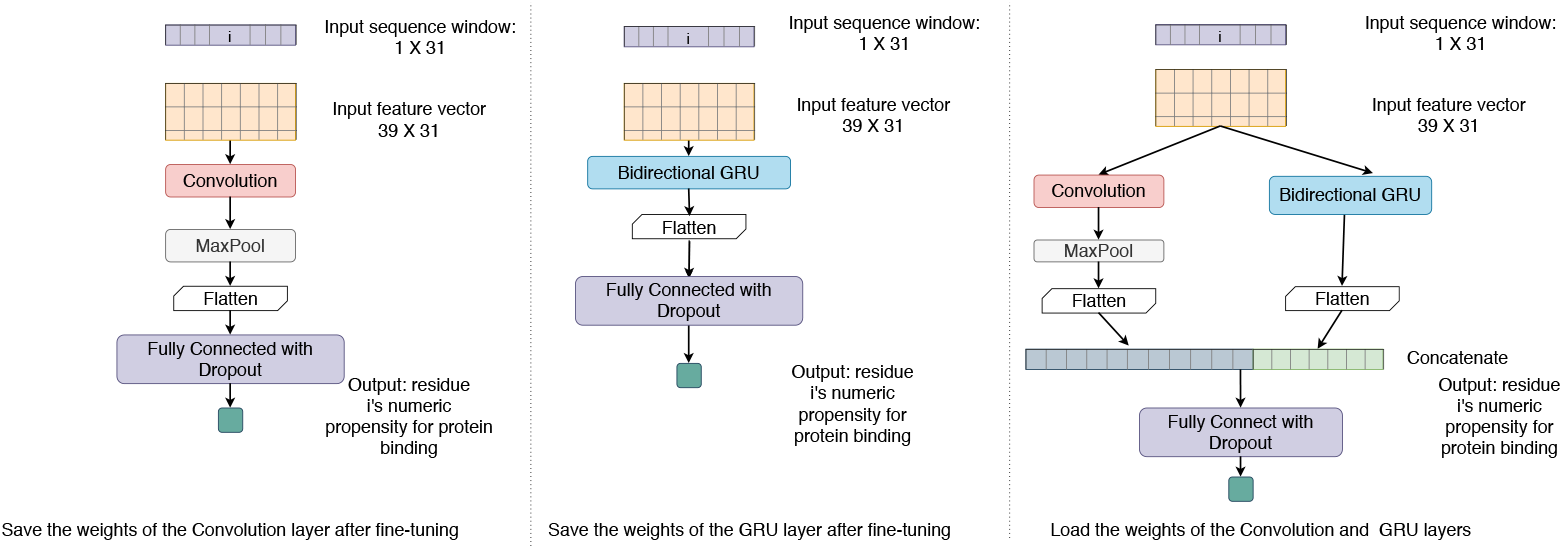
The architecture of DELPHI. Left: the CNN component of the model. Middle: the RNN component of the model. Right: The ensemble model.

Another very useful characteristic of the model is its many-to-one structure, meaning that the information of many residues are used to prediction the binding propensity of the centered single residue. As illustrated in Fig. 2, for each amino acid as the prediction target, a window of size 31, centred on the amino acid position, is used to collect information from the neighbouring 30 residues to help the prediction. A sliding window is used to capture each 31-mer. The size 31 is determined experimentally. The beginning and the ending part of the sequence are padded with zeros. The many-to-one structure has two advantages. Firstly, it serves as a data augmentation technique. Deep learning models need large amount of data to train, and comparing to image classifiers, models in proteomics have access to orders of magnitude less data. Using each residue multiple times during the training process helps the model learn better. Secondly, it makes the model more robust. The lengths of protein sequence vary from less than one hundred to several thousand, and most a many-to-many models have a fixed input length of near 500. During training, sequences around length 500 are often picked. However, during testing, input sequences are random and need to be either padded or cut into pieces. The different average lengths between training and testing could potentially make the model less general.

**Figure 2.**
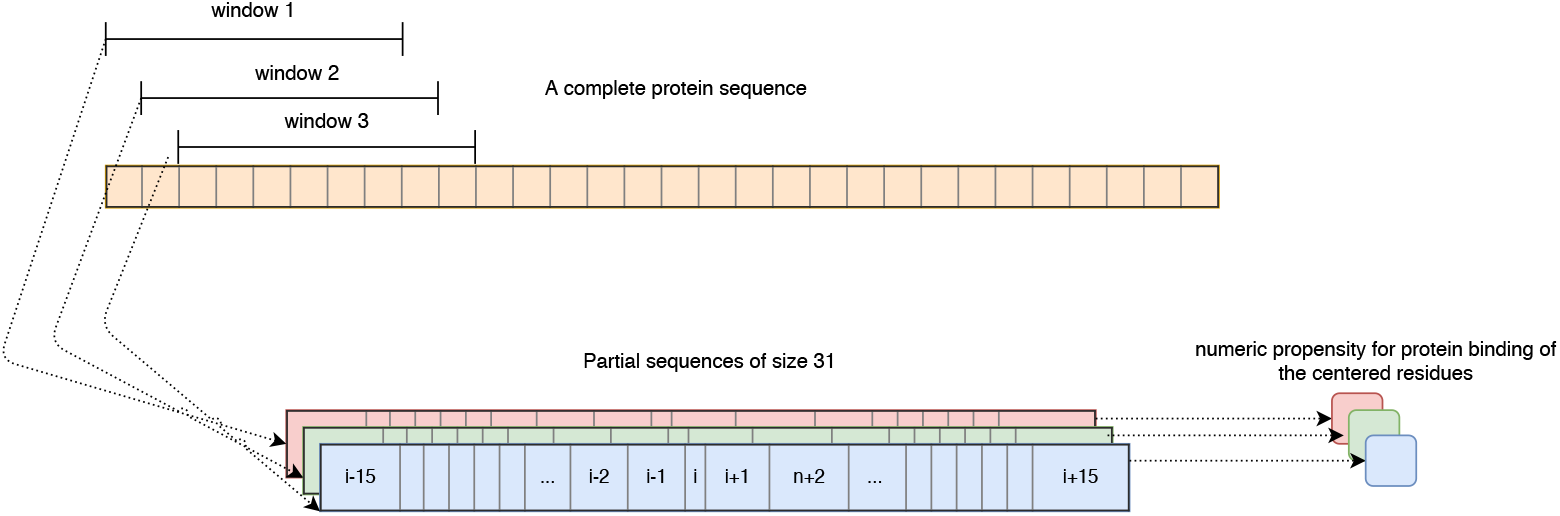
The many-to-one prediction. Sliding windows of size 31, stride 1 are put on top of an input protein sequence. Each time, a sub-sequence of length 31 is extracted. The model predicts the protein-binding propensity of the middle amino acid for each sub-sequence.

### Architecture of the CNN network

The CNN model one has a concise structure: one convolution layer, one maxpooling layer, one flatten layer, and two fully connected layers. For each input sub-sequence of size 31, a 2D feature profile of size 39 by 31 is constructed. The 2D vector is reshaped into 3D and then passed to a convolution 2D layer, followed by a maxpooling layer. The intuition of using the convolution and maxpooling layers is that a 2D protein profile vector can be considered as an image with one channel, and the CNN model captures the combination of several features in a partial image. The results are flattened and then fed into two fully connected layers with dropout for regularization. The last fully connected layer has one unit with activation function sigmoid, so that the output is a single value between 0 to 1. The higher the value, the more confident the CNN model claims that the residue is a PPI binding site.

### Architecture of the RNN network

The RNN component has the following structure: one bidirectional GRU layer, one flatten layer, and two fully connected layers. Similar to the CNN component, a 2D feature profile of size 39 by 31 is built for each 31-mer. The feature profile is passed to a bidirectional gated recurrent units (GRU) layer with the intention to memories the dependency and relationship among the 31 residues. We set the GRU layer to return a whole sequence as opposed to return a single value. The results are flattened and fed into two fully connected layers with dropout. The output of the RNN network is also a single value between 0 to 1.

### Architecture of the ensemble network

The final model combines the core layers of the above mentioned CNN and RNN models and tries to further extract essential information of protein binding. The ensemble network takes a sequence of length 31 as its input. Similar to the CNN and RNN components, a 39 by 31 feature vector is constructed and passed to both a convolution layer and a bidirectional GRU layer. The output of the convolution layer is passed on to a maxpool layer and then flattened. The GRU output is also flattened. Then the outputs of the two flatten layers are concatenated and passed on to two fully connected layers with dropout. The last fully connected layer has one output unit with a sigmoid activation function, so the final output is a single value between 0 to 1, indicating the propensity of being binding sites. This is the final output of the entire model.

Fine tuning is used in this ensemble model. The convolution layer in the CNN network and the bidirectional GRU layer in the RNN network are tuned separated using the same training/validation dataset. After achieving the best performance on the CNN and the RNN components, the weights of the convolution and the GRU layer are saved to files. In the ensemble model, the convolution and the GRU layer load the saved weights from the file and freeze the weights, so that during the process of training, the convolution and the GRU layer stay unchanged. Training and validation data are used again only to train the fully connected layers in the ensemble model.

### Implementation

The program is written in Keras [12] with TensorFlow GPU [1] back end. All features are computed from sequence only. We alleviate the burden of feature computation from users by providing all computation programs and a pipeline script. We ease the system configuration process by providing users a pip package list which enables one-command installation.

Classifying protein binding residue is an imbalanced problem. To cope with that, different class weights [42] are assigned to the positive and negative samples, so that the model pays more attention to the minority class, which is the binding sites. The values are determined by the inverse of the class distribution in the training datasets. In our program, the weights are 0.55 and 4.97 for the non-binding and binding sites respectively.

During training, we shuffle the data before each epoch. Since the sliding window is used to extract each 31-mer, adjacent data entries are very similar; only the first and the last residue differ from the previous and the next data entry. Shuffling the whole training data diversifies the input in each batch. We experimentally trained the model with and without data shuffling, and shuffling the data rendered better predictions.

### Parameter tuning

Parameters and hyper-parameters are chosen based on the training dataset while applying early stopping [37] on the validation set. Early stopping halts the training process when a performance drop on the validation set is detected. This is to avoid overfitting the training dataset. We chose all parameters with the purpose to maximize area under the precision-recall curve (AUPRC) of the training data. All testing results are then carried using the already tuned model. All parameters and hyper-parameters used in this model are shown in Table 3.

**Table 3.** Parameters used in DELPHI. Parameters are divided into four groups: CNN, RNN, ensemble model, and hyper-parameters.

| Parameter | Value |
| --- | --- |
| Epoch in CNN | 8 |
| Kernel size in CNN | 5 |
| Stride in CNN | 1 |
| Padding in CNN | valid |
| Number of filters in CNN | 64 |
| Fully connected unit in CNN | 64, 1 |
| Epoch in RNN | 9 |
| GRU unit | 32 |
| Fully connected unit in RNN | 64, 1 |
| Epoch in ensemble | 5 |
| Fully connected unit in ensemble | 96, 1 |
| Batch size | 1024 |
| Dropout rate | 0.3 |
| Optimizer | Adam ($\beta_1 = 0.9, \beta_2 = 0.999$) |
| Patience in early stop | 4 |
| Loss function | binary cross entropy |
| Learning rate | 0.002 |

The DELPHI model takes 2.1 hours to train the CNN component, 0.5 hour to train the RNN component and 1.3 hours to train the ensemble model on our testing cluster.

## Results

### Competing methods

We have comprehensively compared DELPHI with nine state-of-the-art machine learning based methods. The methods are selected using the following criteria. First, the program is a sequence-based method as sequence information is readily available for most proteins. Second, the program is available in the form of source code or web server. Lastly, the program takes in any input sequence in FASTA format and produces the results on an average-length protein within thirty minutes. Following these criteria, DLpred [52], SCRIBER [54], SSWRF [45], SPRINT [41], CRF-PPI [46], LORIS [13], SPRINGS [40], PSIVER [33], and SPPIDER [36] are selected.

All competing methods are pre-trained using their own training and validation datasets. The most revent two programs, DLPred and SCRIBER, use 5719 and 843 training proteins respectively. The training dataset of DLPred is obtained from CullPDB datasets [?] and further filtered by the authors. The SCRIBER training dataset is originally from the BioLip database. This dataset contains also protein binding information with DNA, RNA, and ligand, which is used by SCRIBER.

All tests have been performed on a Linux (Ubuntu 16.04) machine with 24 CPUs (Intel Xeon v4, 3.00GHz), 256GB memory, and a Nvidia Tesla K40c GPU.

### Evaluation scheme

Similar to previous studies, we use sensitivity, specificity, precision, accuracy, F1-measure, (F1), Matthews correlation coefficient (MCC), area under the receiver operating characteristic curve (AUROC), and area under the precision-recall (AUPRC) to measure the prediction performance. All programs output a prediction value for each amino acid, and thus the receiver operating characteristic (ROC) curve and the precision-recall (PR) curve can be drawn. AUROC and AUPRC are computed based on the curves using Scikit-learn [35]. We focus more on AUROC and AUPR because they are threshold independent and convey an overall performance measurement of a program. The rest of the metrics are calculated using a binding threshold which is determined after obtaining the prediction scores from each program. Since each program’s output is of different scale, for each program, we pick the threshold such that for a given testing dataset, the number of predicted scores above the threshold is equal to the real number of binding sites in the dataset.

The formulas for calculating the metrics are as follows, where true positives (TP) and true negatives (*TN*) are the correctly predicted binding sites and non-binding sites, respectively, and false positives (*FP*) and false negative (*FN*) are incorrectly predicted binding sites and non-binding sites, respectively.

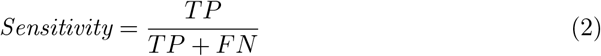

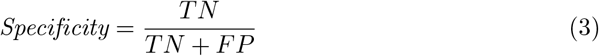

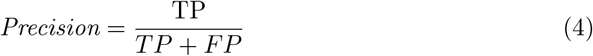

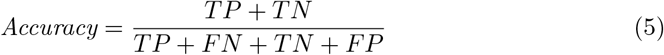

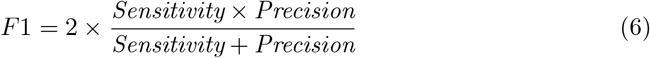

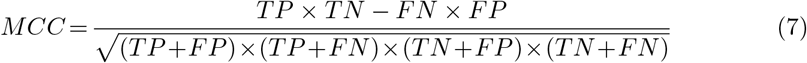

### Comparative assessment of predictive performance

#### Performance comparison on Dset_448

We first compare the DELPHI model with eight programs on Dset_448. This dataset is the most recently published and has the largest number of proteins, so we emphasize the importance of this dataset. As shown in Table 4, DELPHI surpasses competitors in all metrics with an improvement of 3.08% and 17.4% on AUROC and AUPR respectively comparing to the second best program SCRIBER.

**Table 4.** Performance comparison of nine programs on Dset_448. Programs are sorted in asending order by AUPRC. Darker colours represent better results. The evaluation of the programs marked with * are carried by [54].

| Predictor | Sensitivity | Specificity | Precision | Accuracy | F1 | MCC | AUROC | AUPRC |
| --- | --- | --- | --- | --- | --- | --- | --- | --- |
| Dset_448 |  |  |  |  |  |  |  |  |
| SPPIDER* | 0.202 | 0.870 | 0.194 | 0.781 | 0.198 | 0.071 | 0.517 | 0.159 |
| SPRINT* | 0.183 | 0.873 | 0.183 | 0.781 | 0.183 | 0.057 | 0.570 | 0.167 |
| PSIVER* | 0.191 | 0.874 | 0.191 | 0.783 | 0.191 | 0.066 | 0.581 | 0.170 |
| SPRINGS* | 0.229 | 0.882 | 0.228 | 0.796 | 0.229 | 0.111 | 0.625 | 0.201 |
| LORIS* | 0.264 | 0.887 | 0.263 | 0.805 | 0.263 | 0.151 | 0.656 | 0.228 |
| CRFPPI* | 0.268 | 0.887 | 0.264 | 0.805 | 0.266 | 0.154 | 0.681 | 0.238 |
| SSWRF* | 0.288 | 0.891 | 0.286 | 0.811 | 0.287 | 0.178 | 0.687 | 0.256 |
| SCRIBER | 0.334 | 0.896 | 0.332 | 0.821 | 0.333 | 0.230 | 0.715 | 0.287 |
| DELPHI | 0.371 | 0.901 | 0.371 | 0.829 | 0.371 | 0.272 | 0.737 | 0.337 |

In order to have a more comprehensive comparison, we compare DELPHI with another recent sequence-based program, DLPred. However, the training data in DLPred has a big overlap with Dset_448, so we removed the highly similar sequence in Dset_448 using the protocol described in section and produced a reduced dataset Dset_355 which is tested in Section 0.1.

#### Performance comparison on Dset_355, Dset_186, Dset_164, and Dset_72

To further compare DELPHI with other programs, we used Dset_355 along with another three previously published datasets: Dset_186, Dset_164, and Dset_72. Since DLPred and SCRIBER are the most recent programs and have been shown to outperform older programs, we compare DELPHI only with SCRIBER and DLPred on the remaining datasets. As shown in Table 5, DELPHI clearly outperforms the competitors in all metrics on all datasets it shares the least similarities to the testing datasets. As binding site prediction is a highly imbalanced task, the area under the PR curve is a better indication of the performance as it emphasises more on the minority class and ROC curve pays attention to both minority and the majority class [?]. The AUPR is improved by 18.5%, 10.0%, 0.6%, 10.2% comparing to the second best program on each dataset. We present also the average values over Dset_355, Dset_186, Dset_164, and Dset_72 in Table 5 for each specificity value. The performance of DELPHI again surpasses all competing programs. The average AUPR improvement is 9.7%.

**Table 5.**
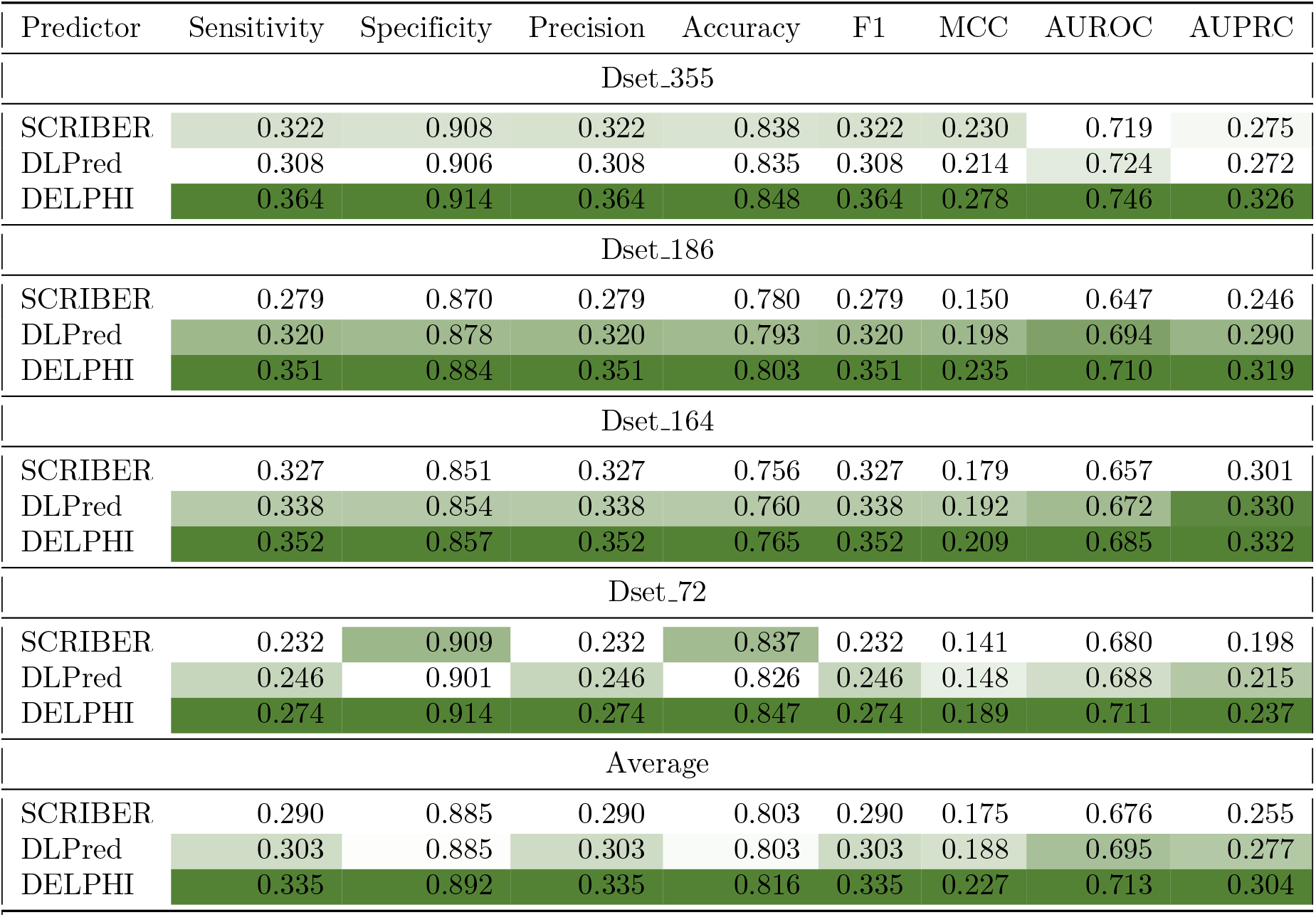
Performance comparison of SCRIBER, DLPred, and DELPHI on Dset_355, Dset_186, Dset_164, and Dset_72. Darker colours represent better results. Each dataset is tested separately using the same metrics. The average metrics of the three datasets is also shown.

| Predictor | Sensitivity | Specificity | Precision | Accuracy | F1 | MCC | AUROC | AUPRC |
| --- | --- | --- | --- | --- | --- | --- | --- | --- |
| Dset_355 |  |  |  |  |  |  |  |  |
| SCRIBER | 0.322 | 0.908 | 0.322 | 0.838 | 0.322 | 0.230 | 0.719 | 0.275 |
| DLPred | 0.308 | 0.906 | 0.308 | 0.835 | 0.308 | 0.214 | 0.724 | 0.272 |
| DELPHI | 0.364 | 0.914 | 0.364 | 0.848 | 0.364 | 0.278 | 0.746 | 0.326 |
| Dset_186 |  |  |  |  |  |  |  |  |
| SCRIBER | 0.279 | 0.870 | 0.279 | 0.780 | 0.279 | 0.150 | 0.647 | 0.246 |
| DLPred | 0.320 | 0.878 | 0.320 | 0.793 | 0.320 | 0.198 | 0.694 | 0.290 |
| DELPHI | 0.351 | 0.884 | 0.351 | 0.803 | 0.351 | 0.235 | 0.710 | 0.319 |
| Dset_164 |  |  |  |  |  |  |  |  |
| SCRIBER | 0.327 | 0.851 | 0.327 | 0.756 | 0.327 | 0.179 | 0.657 | 0.301 |
| DLPred | 0.338 | 0.854 | 0.338 | 0.760 | 0.338 | 0.192 | 0.672 | 0.330 |
| DELPHI | 0.352 | 0.857 | 0.352 | 0.765 | 0.352 | 0.209 | 0.685 | 0.332 |
| Dset_72 |  |  |  |  |  |  |  |  |
| SCRIBER | 0.232 | 0.909 | 0.232 | 0.837 | 0.232 | 0.141 | 0.680 | 0.198 |
| DLPred | 0.246 | 0.901 | 0.246 | 0.826 | 0.246 | 0.148 | 0.688 | 0.215 |
| DELPHI | 0.274 | 0.914 | 0.274 | 0.847 | 0.274 | 0.189 | 0.711 | 0.237 |
| Average |  |  |  |  |  |  |  |  |
| SCRIBER | 0.290 | 0.885 | 0.290 | 0.803 | 0.290 | 0.175 | 0.676 | 0.255 |
| DLPred | 0.303 | 0.885 | 0.303 | 0.803 | 0.303 | 0.188 | 0.695 | 0.277 |
| DELPHI | 0.335 | 0.892 | 0.335 | 0.816 | 0.335 | 0.227 | 0.713 | 0.304 |

#### Feature evaluation

We conducted an another experiment to show that all twelve feature are necessary for DELPHI. We pruned one feature each time, and the remaining eleven features are used to train and then evaluate the DELPHI model. As shown in Fig. 3, the performance decreases with the removal of any feature, showing that there are no redundant features. It is perhaps expected that removing PSSM creates the biggest performance drop, but our newly introduced features, HSP, ProtVec1D, and Position are shown to be very useful as well.

**Figure 3.**
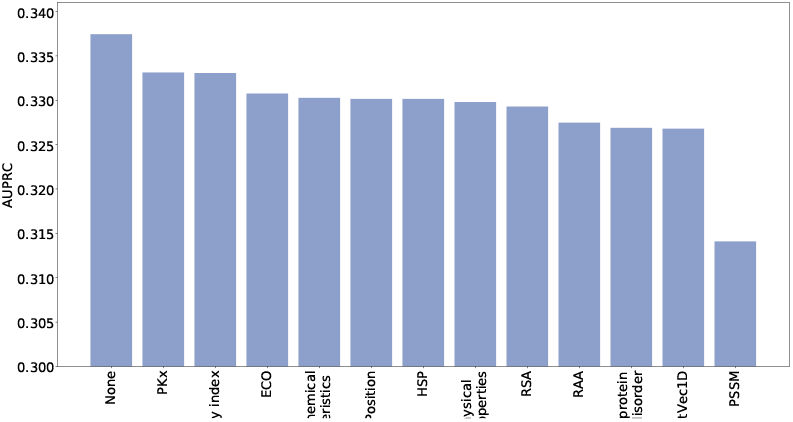
The areas under PR curves with the removal of one out of the twelve features on Dset_448. One feature is removed each time, and the DELPHI model is trained, validated, and tested using the remaining eleven features. The x-axis shows the removed features where ‘None’ indicates using all twelve features, and the y-axis is the AUPR achieved. The features are sorted by the AUPR values.

### Software availability

DELPHI is an open source program under the GPLv3 License. The trained model, source code of training, predicting, and data processing is freely available at https://github.com/lucian-ilie/DELPHI. All datasets used in this study can be downloaded at http://www.csd.uwo.ca/~ilie/DELPHI/.

## Conclusion

We have presented a new deep learning model and program DELPHI, for predicting PPI binding sites. We compared DELPHI with nine current state-of-the-art programs on five datasets and demonstrated that DELPHI has a higher prediction performance. There is still plenty room for improvement on this topic as the highest AUC in all test is 0.747. We hope in the future, the model architecture, the usage of the two new features, and the many-to-one structure can be extended to predicting protein biding with other types of molecules, such as DNA, RNA, and ligand.

## Acknowledgements

We would like to thank Jian Zhang and Buzhong Zhang for running their programs on some of the testing datasets as well as the insightful information they provided regarding SCRIBER and DLPred. We appreciate Giancarlo Colmenares for the suggestions on tuning the network and Min Zeng for the explaination on DeepPPISP.

